# Noisy hemoglobin consumption by erythrocytic *Plasmodium falciparum* parasites: origins and possible implications

**DOI:** 10.1101/2024.09.16.613038

**Authors:** Joseph R. Perko, Abhyudai Singh, Secilia I. Lopez, Shirley Luckhart, Andreas E. Vasdekis

## Abstract

Non-genetic cell-to-cell phenotypic differences can significantly impact pathogen physiology and virulence, leading to unexpected phenomena such as antibiotic persistence. Here, we introduce the role of such non-genetic phenotypic differences in the host, with a focus on hemoglobin consumption by *Plasmodium falciparum* during the erythrocytic stage of parasite development. Through imaging, we quantified the substantial variability in hemoglobin (Hb) concentration among uninfected red blood cells (RBCs), and subsequently measured the rate of Hb consumption by parasites at different stages of their life cycle. This revealed a similarly significant variability among different infected RBCs. By employing a mathematical model, we demonstrated that this variability in Hb consumption can be attributed to non-genetic differences in host RBCs, marking the first evidence of this phenomenon in malaria parasite physiology. These findings underscore the importance of incorporating non-genetic host variability into models of disease progression and treatment strategies for malaria and potentially other pathogen-related diseases.

## Introduction

Non-genetic cell-to-cell phenotypic differences can derive from intrinsic factors (e.g., stochastic fluctuations in gene product levels) and extrinsic factors (e.g., cell age and variations in the extracellular environment).^1–8^ These non-genetic differences can play considerable roles in gene expression and, as such, in cellular stress responses, cell-cycle duration, and metabolic dynamics.^8–14^ By extension, non-genetic phenotypic differences can also influence pathogen virulence. This is particularly true during the initiation of infection that typically involves a small number of infecting organisms.^15^ In this context, investigations of non-genetic phenotypic differences have primarily focused on the pathogen phenotype, leading to several surprising discoveries. These discoveries include the emergence of bet hedging pathogen survival strategies, as well as antibiotic persistence and tolerance.^16–20^

In addition to affecting the pathogen phenotype, non-genetic cell-to-cell differences can also affect the phenotype(s) of host cells that the pathogens infect. Given the complex relationship between host cells and pathogens (e.g., receptor availability in the host, levels of host nutrients)^21–26^, non-genetic phenotypic variations in host cells have the potential to also influence pathogen virulence. Despite this potential, and to the best of our knowledge, the impact of non-genetic host cell-to-cell differences on pathogen biology have not been thoroughly explored. Narrowing this knowledge gap, in conjunction with studies of genetic variability, could lead to more accurate mathematical models of disease progression, as well as breakthroughs in tailoring more effective therapeutic interventions. Here, we specifically explore the influence of non-genetic phenotypic differences in the host cells of erythrocytic-stage *Plasmodium falciparum* malaria *in vitro*.

Malaria is one of three major global infectious diseases, along with HIV/AIDS and tuberculosis, predominantly affecting impoverished populations in sub-Saharan Africa and Southeast Asia.^27^ In these regions, infection with *Plasmodium falciparum* results in the greatest number of human deaths due to malaria.^27^ The *P. falciparum* life cycle begins with transmission via the bite of an infected *Anopheles* spp. mosquito, introducing the parasite into the human bloodstream and initiating a complex developmental process within the host.^28^ Following mosquito transmission, *P. falciparum* parasites enter the human liver, undergo asexual reproduction, and subsequently infect red blood cells (RBCs), resulting in symptomatic malaria. During the erythrocytic stage of the life cycle, parasites consume hemoglobin (Hb) within RBCs, an essential process for parasite virulence as it provides the necessary nutrients for growth and replication.^29^ Notably, Hb consumption is also implicated in the activation of artemisinin (ART)^30, 31^, the frontline treatment for falciparum malaria.^32, 33^

In this report, we investigated the impact of non-genetic host variability on Hb consumption by *P. falciparum*. Employing a single-cell biology approach that relies on optical imaging, we first uncovered significant cell-to-cell differences in the Hb concentration among uninfected host RBCs. We subsequently quantified the rate of Hb consumption by parasites as a function of the parasite life cycle stage. This analysis revealed considerable variability among different infected RBCs. Using a mathematical model, we determined that the variability in Hb consumption can be attributed to host heterogeneity in the Hb concentration of uninfected RBCs. These findings provide first evidence that host cell Hb heterogeneity plays a role in malaria parasite development. Collectively, these observations can support precision models for cell cycle-related disease progression and malaria treatment, particularly for drugs with mechanisms of action that are dependent on Hb degradation, such as ART. This underscores the importance of considering host cell variability in the development of strategies not only for malaria treatment and management, but potentially other infectious diseases as well.

## Results

### Multimodal Imaging

#### Hb quantification

Recognizing the proven effectiveness of Quantitative Phase Imaging (QPI) in quantifying Hb concentration in both infected and uninfected RBCs,^34–37^ we deployed this optical method in this investigation. Generally, QPI operates by measuring the optical path length differences introduced by the specimen at each point in the field of view (**Fig. 1a**). These optical path length differences correlate with the cell’s refractive index, which in turn relates to the cytosolic protein concentration, predominantly comprised of Hb in RBCs.^38^ Thus, QPI is a completely label-free method that eliminates the need to stain infected RBCs for determining Hb concentration, unlike other, recently published methods that do require staining.^39, 40^ Here, we specifically employed the Spatial Light Interferometric Microscopy (SLIM) QPI module, the details of which are detailed elsewhere^41^ and in the **Methods** section. We selected SLIM for its superior 2D imaging capabilities, which allows for the high-throughput screening rates essential to this work. Using QPI measurements, we calculated the Hb concentration within RBCs by first considering the average phase shift of the whole RBC (ΔΦ_ave_) and its direct proportionality to the dry mass density (non-aqueous content) of the cell. For infected RBCs, ΔΦ_ave_ specifically referred to the non-parasitized regions of the cell. Subsequently, we quantified the Hb concentration ([Hb] in g/L) from the measured phase map (ΔΦ_ave_) using the fol-lowing expression^35^ (**Fig. 1b**):

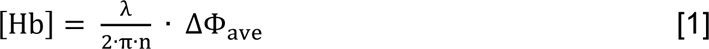

**Fig. 1:**
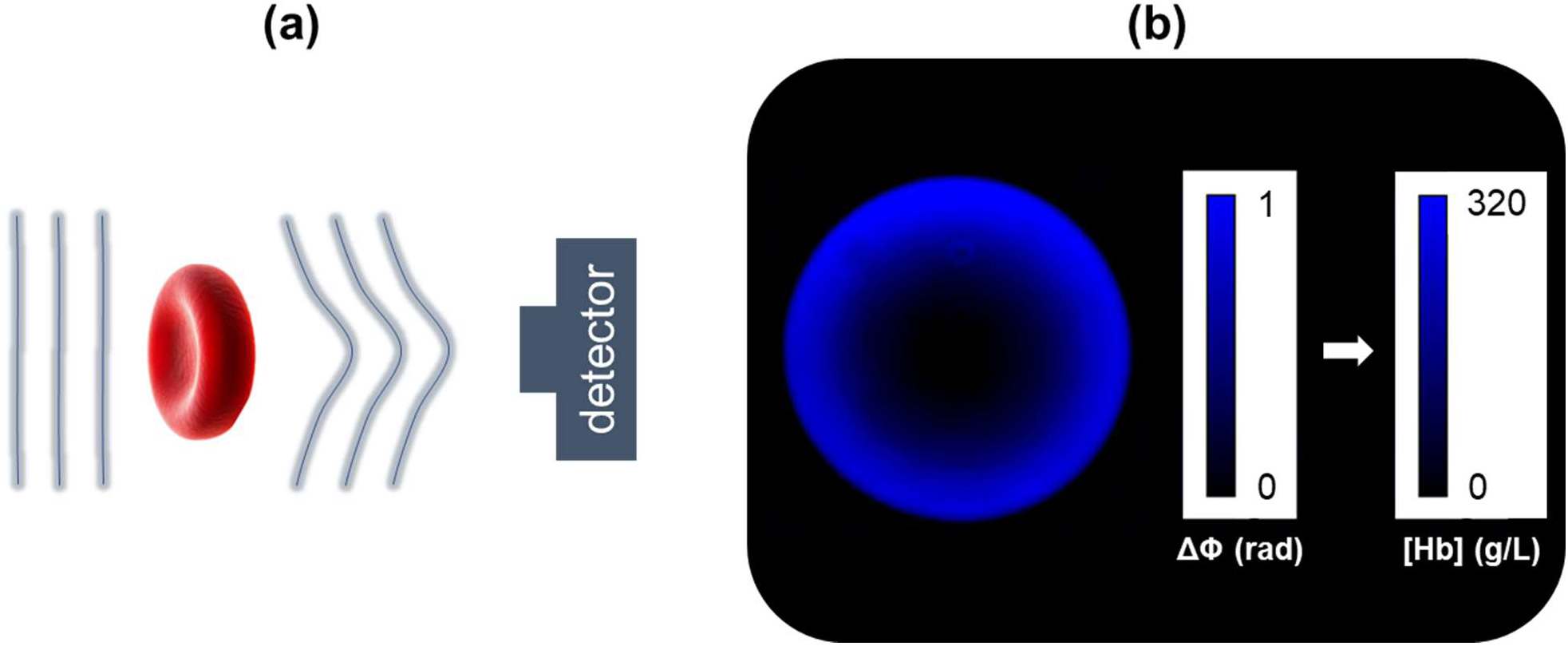
**(a)** A plane wave undergoing wavefront distortion as it transmits through the RBC, which is subsequently recorded in a camera detector; the degree of wavefront distortion depends on the local Hb concentration of the RBC. **(b)** An exemplary QPI image of an RBC; scale bars denote the wavefront distortion (ΔΦ, left) and its conversion to local Hb concentration (in g/L, right).

Where “λ” is the center wavelength of illumination (0.55 μm) and “n” is the specific refractive index increment for Hb (0.2 ml/g).^42^ Through the quantification of Hb concentration and independent measurements of cell size (area), we were also able to quantify the corpuscular Hb per RBC using the previously reported expression:^38^

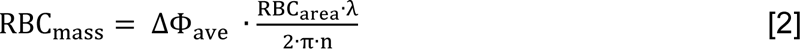

#### Stage classification

In addition to the QPI-based quantification of Hb concentration, we also performed fluorescence (FL) imaging of the same infected RBCs stained with propidium iodide to directly visualize the parasite nucleus and, thus, determine the life cycle stage of the parasite. Unlike recent developments in QPI^43^, the SLIM module^41^ suffers from inevitable optical losses from its polarization-dependent transmission. To minimize the effect of these losses, we acquired the FL images on a different imaging path and a more sensitive imaging sensor (**Methods**). To carry out the stage classification processes, we first cropped the acquired images to 400×400-pixel areas. We then presented the cropped images to two trained users (JRP and SIL), who independently classified parasites into ring (R), late ring or early trophozoite (LR-ET), trophozoite (T), and schizont (Sch). In this context, the parasite life stages were classified as:^44^

- R = ring structure with a thin ring perimeter.
- LR/ET = ring structure with a thicker ring perimeter.
- T = one or two large nuclei, with RBC hosts maintaining their original shape.
- Sch = more than two nuclei within an RBC, but clear merozoites.

**Fig. 2** shows representative images of *P. falciparum* parasites in different life cycle stages from our dataset, with the parasite size increasing from early to late stages. In total, we classified 1,700 RBCs, with parasites in different life stages (R, LR/ET, T, Sch). As further discussed in the **Methods** section, we noted a relatively limited class imbalance in the number of parasites in different life stages. As such, we saw no need for further adjust-ments.

**Fig. 2:**
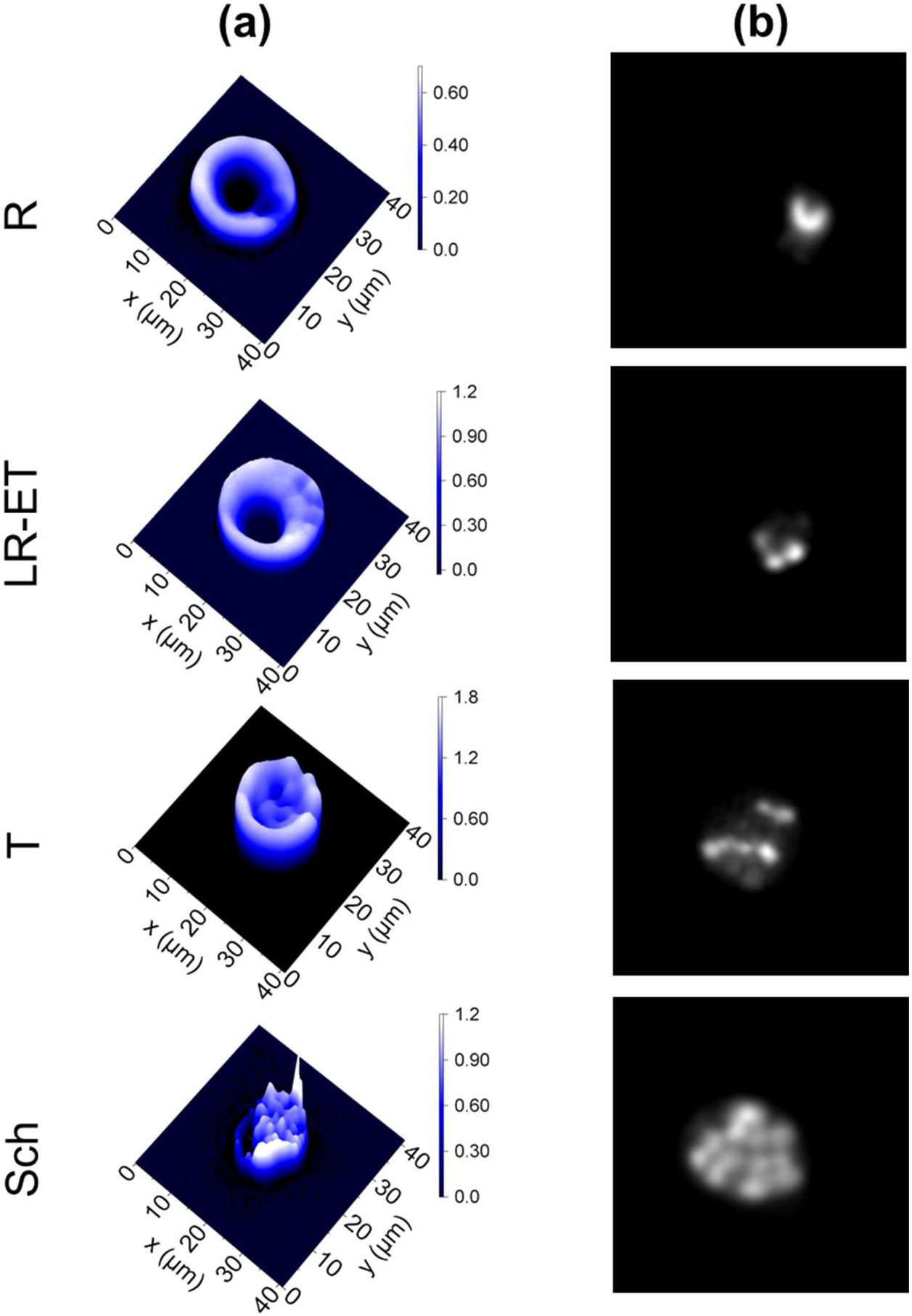
**(a)** QPI and **(b)** fluorescence maps of infected RBCs at different life cycle stages (R: rings; LR-ET: late rings – early trophozoites; T: trophozoites; Sch: schizonts); scale bars in **(b)** denote the wavefront distortion (ΔΦ) in radians.

### Non-genetic host differences

Using our quantitative-phase imaging (QPI) setup, we first imaged uninfected RBCs (**Methods**). Remarkably, we observed substantial cell-to-cell non-genetic differences in cell size, Hb concentration, and consequently, in the corpuscular Hb mass of individual RBCs (**Fig. 3a**). Specifically, we quantified these non-genetic host differences using the coefficient of variation (CoV) metric, which we found to be 21.5% in cell size, 17.1% in Hb concentration, and 26.4% in corpuscular Hb. Similar findings have been reported elsewhere^36^, including notable cell-size differences linked to an optimal RBC size for oxygen advection in arterioles.^45^ Concurrently, comparable levels of non-genetic variability in corpuscular Hb to those observed here have been shown to persist regardless of blood storage conditions.^36^ Interestingly, we observed a weak negative correlation between RBC size and Hb concentration (**Fig. 3b**). However, this correlation was not significant, suggesting that the regulation of RBC size and Hb concentration are likely independent during erythropoiesis. In contrast, we noted a strong positive correlation between cell size and Hb mass^37^ (**Fig. 3c**), which suggests an optimal level for Hb concentration, as earlier reported for RBC size.^45^

**Fig. 3:**
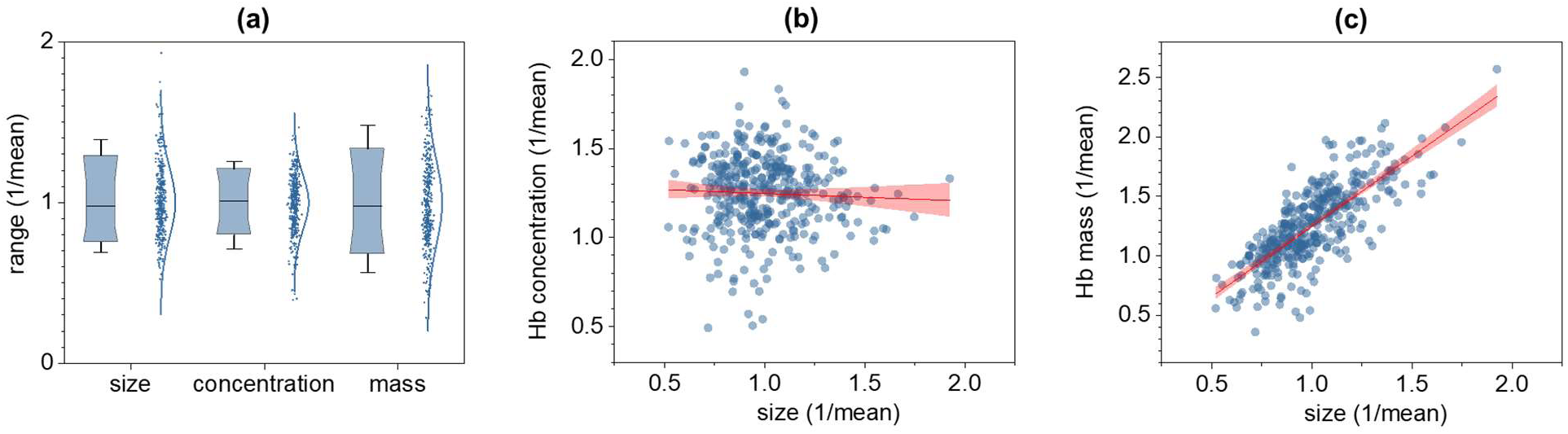
**(a)** RBC cell-to-cell differences in size, Hb concentration, and corpuscular Hb (all normalized over the mean); box charts represent 10%, 25%, 70%, and 90% ranges, while whiskers represent the 5%-95% range. **(b)** The Hb concentration of uninfected RBCs normalized to the population mean as a function of the RBC cell size (also normalized over the mean); red line represents the linear regression analysis and its 95% confidence intervals (red shaded area); the slope was determined by bootstrapping (10,000 samples, **Methods**) and was found to be −0.0341 with [−0.1030, 0.0381] 95% confidence intervals. **(c)** Same as in **(a)**, albeit for RBC cell size and mass with a slope of 0.9470 and [0.8719, 1.0234] 95% confidence intervals.

### Non-genetic differences in Hb consumption

We then investigated how the Hb concentration of infected erythrocytes changes as a function of the parasite’s life cycle stage. As depicted in **Fig. 4a**, we observed an overall decreasing trend in Hb concentration with advancing parasite life cycle stages, consistent with the parasite consumption of intraerythrocytic Hb.^34^ Notably, Hb concentration varied significantly between infected RBCs, even among those harboring parasites at identical life cycle stages (**Fig. 4a**). Host-to-host cell differences in Hb concentration (quantified through CoV) were 17.1±1.2% for uninfected RBCs, increasing to 31.3±8.4% for infected RBCs with parasites at the ring life cycle stage (**Fig. 4b**). Overall, cell-to-cell Hb concentration differences, or noise were consistently higher in infected RBCs compared to uninfected ones. This observation suggests varying rates of Hb consumption among individual parasites, which could, in turn, predict differences in parasite development rates that are assumed in existing mathematical models of the life cycle distribution of ABS parasites. The notion of differential rates of Hb consumption between individual parasites is further corroborated by the abrupt increase in cell-to-cell variability in Hb consumption at the schizont stage (**Fig. 4b**), where Hb consumption is also intensified (**Fig. 4a**). With these observations in mind, a key question arises: what potential mechanisms could be driving the observed variations in Hb consumption? To address this question, we undertook mathematical modeling and subsequent fitting to the experimental data (**Fig. 4**). Our model was inspired by Lotka-Volterra predator-prey dynamics and adapted to describe the dynamic interaction between Hb concentration and parasite biomass (**Methods**).^46, 47^ By fitting our model to the experimental data, we inferred critical parameters that describe the rate of Hb consumption and its conversion into parasite biomass. To capture the transient variability in Hb decay, we considered two main sources of heterogeneity: differences in Hb concentration between uninfected RBCs and the grouping of erythrocytic parasites at different life cycle stages. In essence, the first source reflects the non-genetic cell-to-cell variability in host RBCs, while the second acknowledges the uncertainty in the exact time each parasite spends in a specific stage prior to imaging. By incorporating these sources of heterogeneity, our model uniquely accounts for host heterogeneity, unmasking a key role in parasite development and highlighting the complexity of Hb dynamics within infected RBCs. This underscores the importance of considering both biological and temporal factors in predicting parasite behavior and eventually treatment outcomes.

**Fig. 4:**
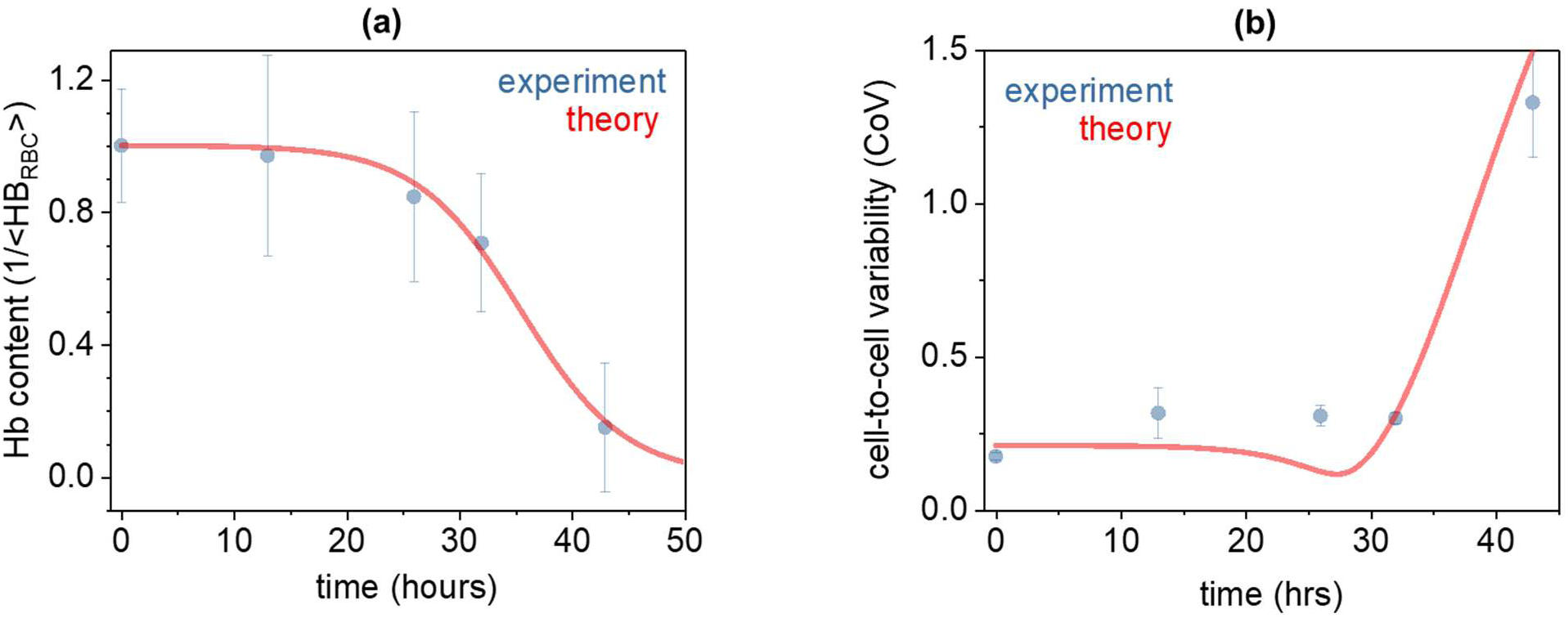
**(a)** The dependence of the Hb concentration on asexual *P. falciparum* developmental stage (normalized to the Hb concentration of the uninfected RBC); datapoints represent the mean ± standard deviation, and the thick red line represents the theoretical result. **(b)** Cell-to-cell Hb concentration differences (quantified through the coefficient of variation, CoV) as a function of the *P. falciparum* developmental stage; points represent experimental CoV data with error bars computed by bootstrapping (10,000 samples, **Methods**); thick red line represents the theoretical result.

## Discussion

In summary, we quantified the non-genetic differences in Hb consumption among erythrocytic *P. falciparum* parasites. Our observations and model suggest that non-genetic cell-to-cell variations in Hb levels among uninfected host RBCs can contribute to the differences in the rates of Hb consumption between parasites that could – by extension – also influence parasite development rates.

In greater detail, we first quantified the cell-to-cell differences in Hb concentration among uninfected RBCs, which yielded a 17% CoV and a four-fold range (i.e., maximum/minimum) in Hb levels (**Fig. 3a**). While Hb concentration did not correlate with cell size (**Fig. 3b**), previous calculations suggest that the observed four-fold range in Hb concentration (or crowding) is significant and can influence the activity coefficient of Hb by an exorbitant 100×.^48^ To better understand the origins of this Hb concentration variability among RBCs, it would be useful to consider the process of RBC generation (erythropoiesis) in the bone marrow.^49^ During erythropoiesis, particularly in the polychromatic and orthochromatic stages,^50^ erythroblasts increase Hb synthesis by upregulating globin gene expression.^51^ This process likely continues through enucleation, though dependent on existing ribosomes since globin gene transcription has ceased.^52^ Therefore, non-genetic cell-to-cell differences in Hb concentration among RBCs could potentially arise from variability in globin gene expression during erythropoiesis, under the influence of cellular noise and stochastic fluctuations.^1–8^ Another possible source of Hb variability between RBCs could be the loss of cytoplasmic material—such as Hb molecules, ribosomes, and globin mRNA—during enucleation.^53^ While experimentally investigating these origins is presently challenging, emerging methodological breakthroughs are paving the way for such granular level explorations.^54^

We also investigated the variability of Hb concentration in infected RBCs across different life cycle stages. Our analysis revealed significant differences in Hb concentration among infected RBCs, even when parasites were at identical stages of development (**Fig. 3b**). Notably, these variations were more pronounced in infected RBCs compared to uninfected ones, suggesting differential rates of Hb consumption by individual parasites (**Fig. 3b**). To confirm this notion, we deployed a mathematical model (**Fig. 4**) and determined that the transient variability in Hb decay stems from both fluctuations in the Hb concentration of uninfected RBCs and the grouping of different time points into distinct parasite stages. This study provides the first insight into how non-genetic differences in host cells could influence parasite physiology. These findings can be extended to explore the known cell-to-cell variability in parasite life cycle duration and developmental rates, as well as the complexity of Hb dynamics within infected RBCs.^55–57^ While this variability has been previously attributed to epigenetic plasticity resulting from chromatin reorganization and frequent mutations, our findings introduce a previously unexplored factor: the non-genetic differences in the Hb concentration of uninfected RBCs.^55–57^ This new understanding of Hb variability could better define the host cell conditions that support parasite progression through various life cycle stages. Future research could explore how these non-genetic differences in Hb consumption impact the efficacy of treatments like ART, and extend this investigation to other pathogens as well.

## Acknowledgments

AEV gratefully acknowledges funding from the U.S. National Science Foundation project 2041523 for supporting JRP and SIL, and the U.S. Department of Energy project DE-SC0022282 for hardware support. SL acknowledges support from the U.S. National Institutes of Health, National Institute for Allergy and Infectious Diseases.

## Contributions

JRP performed experiments and parasite classification. AS performed mathematical modeling. SIL performed experiments and parasite clarification. SL provided biological materials. AEV overviewed the research and wrote the manuscript.

## Methods

### Microscopy

Images were obtained using a quantitative phase imaging system (Phi Optics), which relies on the spatial light interference microscopy (SLIM) principle.^41^ This technique involves creating phase images by projecting the back focal plane of a phase contrast objective onto a liquid crystal phase modulator. This modulator alters the optical phase of the light wavefront scattered by the sample in relation to the un-scattered light.^41^ This method allowed us to capture images showing the relative phase delay between the cells and their cytosolic components (scattered wavefront) compared to the background (un-scattered wavefront).

The QPI system was integrated with an inverted microscope (DMi8, Leica) featuring phase contrast and fluorescent imaging capabilities, an automated XYZ stage, and a 63× magnification objective (NA 0.7, PH2, Leica). Images were captured using the Orca Flash 4.0 scientific CMOS camera (Hamamatsu) with a 6.5 µm pixel size. Acquisition settings included a 50 ms exposure time and a 50 ms refresh rate for the spatial light modulator. This setup provided an optimal balance between stability and temporal resolution, consistent with our previous findings.^58^ For each experimental condition, 2D images were collected by scanning the stage in the XY plane until 1700 single-cell observations were obtained. In our analysis of parasite counts across different life stages, we observed a slight imbalance in the distribution, with each life stage accounting for 20% ± 5% of the observations (mean ± SEM). Given this modest variation, we determined that adjusting for class imbalance was unnecessary.

### Mathematical modeling of Hb consumption

To investigate the mechanisms driving the observed variations in Hb consumption among infected RBCs, we developed a mathematical model influenced by Lotka-Volterra preda-tor-prey dynamics.^46, 47^ This model captures the interaction between Hb concentration and parasite biomass over time. The interaction between Hb concentration (y) and parasite biomass (x) is described by the following nonlinear system of ordinary differential equations:

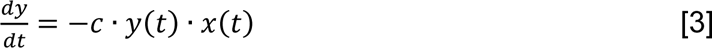

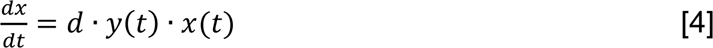

where *y*(*t*), *x*(*t*) are Hb concentration and parasite biomass, at time *t*, respectively. Here *c* denotes the per capita consumption rate of Hb, and the ratio *d*/*c* can be interpreted as the conversion of Hb into parasite biomass. This system has the following solution for Hb levels over time:

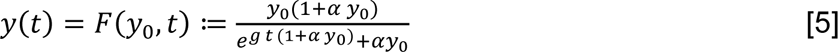

where *α*, *g* are two lumped parameters given by

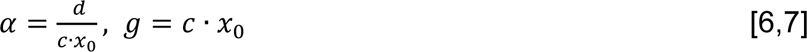

Here *y*_0_ & *x*_0_ are values of *y*(*t*) & *x*(*t*), respectively, at time *t* = 0. Fitting eq. (1) to the data on the decay in average Hb levels over time results in the inference of parameters *g* ≈ 10^−4^ *day*^−l^ and the dimensionless ratio

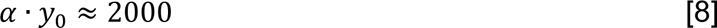

For this fitting, we assumed parasite stage R, LR-ET, T, and S correspond to 13, 26, 32, and 43 hours, respectively.^59^ To capture the transient variability in Hb decay, we considered heterogeneity arising from two sources: fluctuations in initial Hb amount (y_0_) and the lumping of different time points into specific stages. In the limit of small fluctuations in these quantities, the variability in Hb levels (as determined by its coefficient of variation) is given by:

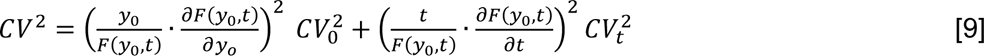

where *F*(*y*_0_, *t*) is given by (1), and from fitting we obtain *CV*_0_ = *CV_t_* = 0.3 that quantify the variability in initial Hb levels, and variability arising as result of temporal clustering of data, respectively.

### Samples

#### Human blood preparation

Human RBCs were prepared from commercially purchased O+ human blood (Grifols Bio Supplies, Inc., Los Angeles, CA, USA) immediately after receipt or, at the latest, the following day, and stored at 4°C. After receipt, blood was transferred into sterile 50 ml conical tubes and supplemented with RPMI 1640 (without HEPES) to 50% vol:vol. The contents were mixed by gently inverting the tubes, followed by centrifugation at 4°C for 10 minutes at 1000g. After centrifugation, the supernatant was aspirated using a glass pipette, with these steps repeated three times to remove white blood cells (WBCs) and proteins. Washed RBCs were transferred to a fresh 50 ml conical tube, mixed with an equal volume of regular RPMI 1640, and stored at 4°C for a maximum of 2 weeks. Washed RBCs were sampled, fixed in 2% paraformaldehyde (overnight at 4°C) and washed (3×) in fresh RPMI 1640.

#### Plasmodium falciparum culture

To initiate the culture, frozen stock of *P. falciparum* NF54-infected RBCs was warmed to 37°C and washed three times with decreasing concentrations of NaCl (12%, 1.6%, 0.9%) to remove glycerol. Washed RBCs were then transferred to small flasks.^60^ Live parasites were grown at 37°C in complete RPMI-1640 medium supplemented with 10% heat-inactivated human serum, HEPES, L-glutamine, hypoxanthine and DL-Lactic acid with 4.0-6.0% washed type O+ RBCs prepared as described above. Medium was changed each day, followed by flow of mixed gas (5% CO_2_, 5% O_2_, 90% N_2_) into the flask. Every two days, the percentage of infected RBCs was determined using microscopic examination of thin blood films stained with Giemsa.^61^ At 10-12% infected RBCs, the parasite culture was split into a new flask for maintenance as above. Infected RBCs for our studies were sampled from these flasks, fixed in 2% paraformaldehyde (overnight at 4°C) and washed three times in fresh medium.

#### Statistics

The coefficient of variation (CoV) was computed in Origin Pro. The 95% confidence intervals of all linear regressions were computed in MATLAB by bootstrapping using the bootci function (n = 10,000 samples).

